# Fractal Dimension of Cortical Functional Connectivity Networks Predicts Severity in Disorders of Consciousness

**DOI:** 10.1101/789636

**Authors:** TF. Varley, M. Craig, R. Adapa, P. Finoia, G. Williams, J. Alanson, J. Pickard, DK. Menon, EA. Stamatakis

## Abstract

Recent evidence suggests that the quantity and quality of conscious experience may be a function of the complexity of activity in the brain, and that consciousness emerges in a critical zone on the axes of order/randomness and integration/differentiation. We propose fractal shapes as a measure of proximity to this critical point, as fractal dimension encodes information about complexity beyond simple entropy or randomness, and fractal structures are known to emerge in systems nearing a critical point. To validate this, we tested the several measures of fractal dimension on the brain activity from healthy volunteers and patients with disorders of consciousness of varying severity. We used a Compact Box Burning algorithm to compute the fractal dimension of cortical functional connectivity networks as well as computing the fractal dimension of the associated adjacency matrices using a 2D box-counting algorithm. To test whether brain activity is fractal in time as well as space, we used the Higuchi temporal fractal dimension on BOLD time-series. We found significant decreases in the fractal dimension between healthy volunteers (n=15), patients in a minimally conscious state (n=10), and patients in a vegetative state (n=8), regardless of the mechanism of injury. We also found significant decreases in adjacency matrix fractal dimension and Higuchi temporal fractal dimension, which correlated with decreasing level of consciousness. These results suggest that cortical functional connectivity networks display fractal character and that this is predictive of level of consciousness in a clinically relevant population, with more fractal (i.e. more complex) networks being associated with higher levels of consciousness. This supports the hypothesis that level of consciousness and system complexity are positively associated, and is consistent with previous EEG, MEG, and fMRI studies.

## 1 Introduction

Research into the neural origins of conscious experience has suggested that consciousness may be associated with the complexity of information integration in the brain (Tononi, 2008; Koch et al., 2016). While complexity, like consciousness, is a difficult thing to define, several measures have emerged in complex systems science for describing what it means for a system to be complex in domains such as system architecture, spatial, and temporal dynamics (Mitchell, 2009). In the context of biology, complexity can refer to how the components of a naturally occurring system interact and encode information, with a particular interest in emergent properties and self-organizing behaviour (Baianu et al., 2007). The brain is often considered to be a paradigmatic example of a complex system, showing many hallmarks of complexity, such as billions of interacting neurons, which encode information, respond to stimuli, compute information, and give rise to astonishing emergent phenomena, the most mysterious of which is consciousness.

The Entropic Brain Hypothesis (EBH) posits that consciousness emerges when the brain is near a critical point between order and randomness, known as criticality, and that to move too far in either direction will result in a change in the quality of consciousness, and ultimately, loss of consciousness entirely (Carhart-Harris et al., 2014; Carhart-Harris, 2018). Various studies have used different metrics to approximate the complexity of brain activity, and the results have been quite consistent, even across modalities. Studies that estimate the Lempel-Ziv complexity of EEG and MEG signals have found that the algorithmic complexity of time-series is decreased in both healthy volunteers and patients who have had their level of consciousness reduced by a range of mechanisms, including sleep (**?**), sedation with anaesthetics (Schartner et al., 2015), and brain injury (Sarasso et al., 2014). Conversely, the complexity of brain signals is increased in volunteers who are under the influence of psychedelic drugs like LSD, suggesting a corresponding increase in the complexity of brain activity (Schartner et al., 2017). Studies have also found that alteration to consciousness is associated with differences in the complexity of functional connectivity networks (Tagliazucchi et al., 2014; Tagliazucchi et al., 2016; Atasoy et al., 2017; Viol et al., 2017; Pappas et al., 2018), which may imply that the spatial complexity of brain activity is as important for the maintenance of consciousness as temporal complexity indexed by MEG and EEG measures.

A core feature of the EBH is that consciousness emerges, not where algorithmic complexity is maximal, but in the critical zone, on the boarder between low- and high-entropy states (“the edge of chaos”). Several publications suggest that the healthy brain operates at, or just below, this area of criticality (Beggs and Timme, 2012; Cocchi et al., 2017), and there are compelling theoretical reasons to prefer a critical model of the brain: in neural networks, critical systems show the greatest ability to perform computations (Shew et al., 2011), store information (Yang et al., 2012), and criticality maximizes the range of input scales (dynamic range), due to the scale free nature of critical activity (Shew et al., 2009). These are all qualities that a brain capable of supporting a complex phenomenon like consciousness might be expected to show. While criticality and complexity, as formalized by theories such as Integrated Information Theory (Tononi, 2008) are separate; *in vivo* and *in silico* studies have found that they are locally maximal in the same regions (Timme et al., 2016). The specifics of criticality in the brain have not gone unchallenged (Kanders et al., 2017), however, studies investigating the relationship between criticality, consciousness, and brain function in disorders of consciousness may shed further light on the topic.

One of the “fingerprints” of critical phenomena is the emergence of scale-free, or fractal behaviour near the critical point (Beggs and Timme, 2012). While the use of fractal analysis on neural structures and signals has been significant (Ieva et al., 2014; Ieva et al., 2015), there has been far less analysis of functional connectivity networks. Gallos et al., (Gallos et al., 2012a; Gallos et al., 2012b) showed that there was a relationship between the fractal structure of a voxel-level functional connectivity network and the threshold at which the edges were removed: if only the strongest connections are kept, the network has pronounced fractal character, however, when weaker connections are incorporated, the fractal character is reduced and replaced by a small-world character. It was further shown that these weak small-world connections provide near optimal integration of information flow between the strong fractal modules. Gallos et al., proposed that this may be a possible solution to the problem of integration versus modularity: the brain processes many different sensory modalities separately, however, the unified nature of conscious experience requires high-level integration of sensory and cognitive processes after processing has occurred separately in primary sensory areas. It may be that functional connectivity networks can be thought of as being divided into two layers: the ‘foundational layer,’ which is made up of the strongest connections forms a large world, fractal backbone that is modular, but not particularly well-integrated, where low-level sensory processing might occur before being bound together by higher, connected layers. The second layer, the ‘integration layer’ is the set of weaker edges that connect the provide integration for the different modules of the ‘foundational layer’ (Gallos et al., 2012a; Gallos et al., 2012b).

Based on these results, we made two hypotheses: (1) functional connectivity networks in the neocortex would have a measurable fractal character when all of the weaker edges had been thresholded out and (2) that in patients with reduced level of consciousness, this fractal character would be degraded. To investigate this, we used resting-state fMRI data from healthy volunteers and patients suffering from reduced levels of consciousness associated with brain injury. We divided these patients into two subgroups based on clinical diagnosis: minimally-conscious state (MCS) and vegetative state (VS), based on accepted diagnostic criteria (Laureys et al., 2004). In general, a higher fractal dimension is associated with a more complex system (Corbit and Garbary, 1995), and so a decrease in fractal dimension in patients with reduced level of consciousness would suggest that the neural activity in those subjects was reduced, which would be consistent with the predictions of the Entropic Brain Hypothesis. To supplement this data, we also used a commonly-used measure of time-series fractal dimension, the Higuchi temporal fractal dimension algorithm (Higuchi, 1988; Klonowski et al., 2010) to determine whether the fractal character of temporal activity follows the same pattern as spatial activity. In EEG studies, the Higuchi fractal dimension has been found to drop when consciousness is lost in sleep (Klonowski et al., 2005) and anaesthesia (Spasic et al., 2011), so we hypothesized a similar effect would be seen in BOLD time-series.

## 2 Materials & Methods

### 2.1 Calculating Network Fractal Dimension

Since the fractal dimension of most real-world systems cannot be solved analytically, researchers commonly use a family of algorithms known as box-counting measures to determine the fractal dimension of a natural system. The box-counting dimension describes how the topology of a surface changes (or remains the same) at different scales. For any shape, two values are defined: *l*_*B*_ which is the length of an *n*-dimensional box and *N* (*l*_*B*_), which is the minimum number of boxes necessary to ‘tile’ the surface in question. the shape being tiled is a fractal, then:

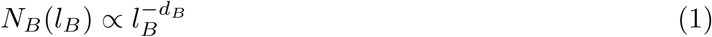

Where *d*_*B*_ is the box-counting dimension. Algebraic manipulation shows that *d*_*B*_ can be extracted by linear regression as:

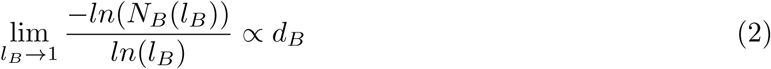

A similar logic is used when calculating the box-counting dimension of a graph. For a graph *G* = (*V, E*), a box with diameter *l*_*B*_ defines a set of nodes *B ⊂ V* where for every pair of nodes *v*_*i*_ and *v*_*j*_ the distance between them *l*_*ij*_ *< l_B_*. To quantify the fractal dimension of the functional connectivity networks, a box counting method, the Compact Box Burning (CBB) algorithm, was used to find *N*_*B*_(*l*_*B*_) for a range of integer *l*_*B*_ values 1..10. If *G* has fractal character, a plot of *ln*(*N*_*B*_(*l*_*B*_)) vs. *ln*(*l*_*B*_) should be roughly linear, with a slope of *-d_B_*. Due to the logarithmic relationship between box-size and fractal dimension, exponentially higher resolutions (in this case, numbers of nodes) are required to achieve modest increases in the accuracy of the measured fractal dimension. Computational explorations, where a box-counting method is used to approximate a fractal dimension that has already been solved analytically, show that the box-counting dimension converges to the true dimension with excruciating slowness (Joosten et al., 2016), necessitating largest network that is computationally tractable. In this context, where each node in our network maps to a specific brain region, we had to segment (parcellate) the cortex into as many distinct brain regions as we could, in this case, using a parcellation with 1000 ROIs (Schaefer et al., 2017).

We need to note that we are not doing a truly rigorous power-law inference. The question of when an empirical distribution can be considered to follow a power-law is a rich field of research (Clauset et al., 2009; Voitalov et al., 2018; Gerlach and Altmann, 2019). To do a statistically rigorous powerlaw inference typically requires multiple decades of values to do a maximum likelihood estimate. Due to the inherent limitations of this box-counting algorithm, such a wide range was impossible. Consequently, we made no strong claims about how well any condition adheres to a power law, but rather, are interested in how multi-scale structure changes between conditions.

#### 2.1.1 FracLac Adjacency Matrix Analysis

Our second test of fractal structure used a two-dimensional box-counting method to analyse the associated adjacency matrix representations of the functional brain networks. This analysis served two purposes: primarily, it was meant to replicate the results of the CBB analysis, however we also hoped that, if it did replicate the initial results, it could be a more computationally efficient method for estimating the fractal dimension of a brain network. By using a different representation of the network, we hoped to show that the quality of network fractal dimension is conserved across isomorphic representations. This would increase our confidence in the CBB results by showing that our findings are unlikely to be an artefact of that particular algorithm. For a given graph *G* = (*V, E*) with nodes *v*_*i*_ and *v*_*j*_, the corresponding adjacency matrix, *A*(*G*) is defined:

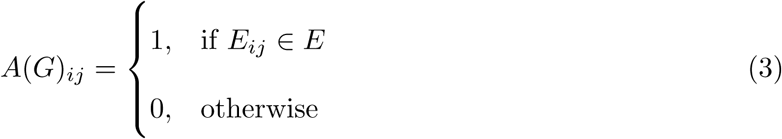

In the resulting matrix, every 1 represents an edge between two nodes *v*_*i*_ and *v*_*j*_. If the distribution of edges *E* ∈*G* is fractal, we hypothesized that the distribution of 1’s in the associated matrix *A*(*G*) would also have fractal character. To test this, we used the program FracLac (Karperien, A., FracLac for ImageJ, version 2015sep09)^1^, a plugin for ImageJ software (Wayne Rasband, National Institutes of Health, USA). FracLac uses a simple, 2-dimensional box-counting algorithm to return the fractal dimension of the distribution of pixels in the image. FracLac returns an upper and lower bound on the range of the fractal dimension for each image, based on the instantaneous value of *d*_*B*_ at every value of *l*_*B*_. For the purposes of this analysis, we took the mean of those values and defined that average as the fractal dimension of each image. The adjacency matrices were exported as binary .jpg images for analysis, and the default values for FracLac’s batch image analysis were used.

We hypothesized that this method, while more accessible and less abstract than the Compact-Box-Burning algorithm, would be less sensitive to small changes in fractal dimension between conditions, as some information is lost when doing a two-dimensional box counting algorithm on a flat network representation, rather than operating directly on a graph.

### 2.2 Higuchi Temporal Fractal Dimension

We used the Higuchi temporal fractal dimension algorithm, widely used in EEG and MEG analysis, to calculate the fractal dimension of temporal brain activity (Higuchi, 1988; Kesi and Spasi, 2016). We will briefly describe the method here. The algorithm takes in a time-series *X*(*t*) with *N* individual samples corresponding to one Hilbert-transformed BOLD time-series extracted from our functional brain scans (details below). From each time-series *X*(*t*), we create a new time-series 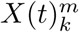, defined as follows:

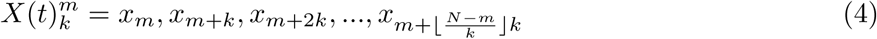

where *m* = 1, 2, …, *k*.

For each time-series 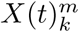 in *k*_1_, *k*_2_, …*k*_*max*_, the length of that series, *L*_*m*_(*k*), is given by:

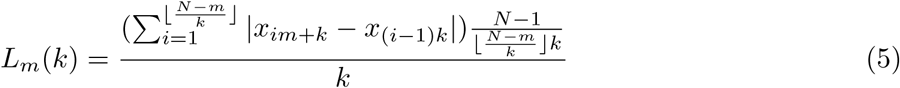

We then define the average length of the series ⟨*L*(*k*)⟩, on the interval [*k, L*_*m*_(*k*)] as:

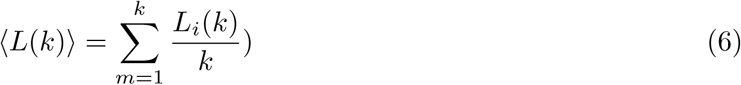

If our initial time-series *X*(*t*) has fractal character, then ⟨*L*(*k*) ⟩ ∝ *k*^*-D*^. As with the procedure for calculating the network fractal dimension, the algorithm iterates through values of *k* from 1…*k*_*max*_ and calculates *ln*(⟨*L*(*k*) ⟩) vs. *ln*(*k*^*-*1^), extracting *D* by linear regression. The various values of *k* can be thought of as analogous to the various values of *l*_*B*_ used to calculate the network fractal dimension. The Higuchi algorithm requires a pre-defined *k*_*max*_ value as an input, along with the target time-series. This value is usually determined by sampling the results returned by different values of *k*_*max*_ and selecting a value based on the range of *k*_*max*_ where the fractal dimension is stable. For both DOC datasets, we chose *k*_*max*_ = 64 as this was the largest value that our algorithm could handle.

The implementation we used was from the PyEEG toolbox (Bao et al., 2011), downloaded from the Anaconda repository.

### 2.3 fMRI Data Acquisition & Preprocessing

#### 2.3.1 Healthy Control Data

Ethical approval for these studies was obtained from the Cambridgeshire 2 Regional Ethics Committee, and all subjects gave informed consent to participate in the study. Twenty five healthy volunteer subjects were recruited for scanning. The acquisition procedures are described in detail in (Stamatakis et al., 2010; Adapa et al., 2014): MRI data were acquired on a Siemens Trio 3T scanner (Wolfson Brain Imaging Center, Cambridge). T1-weighted were acquired using an MP-RAGE sequence (TR = 2250 ms, TI = 900 ms, TE = 2.99 ms and flip angle = 9*°*), with an structural images at 1 mm isotropic resolution in the sagittal plane. Each functional BOLD volume consisted of 32 interleaved, descending, oblique axial slices, 3 mm thick with interslice gap of 0.75 mm and in-plane resolution of 3 mm, field of view = 192 × 192 mm, TR = 2 s, TE = 30 ms, and flip angle 78 deg.

Of the 25 healthy subjects datasets, 10 were excluded, either because of missing scans (n=2), or due of excessive motion in the scanner (n=8, 5mm maximum motion threshold). For this study, we only used the awake, control condition described in the original Stamatakis study, ignoring the drug conditions.

The resulting images were preprocessed using the CONN functional connectivity toolbox (Whitfield-Gabrieli and Nieto-Castanon, 2012), which uses Statistical Parametric Mapping 12 ^2^ and MATLAB version 2017a^3^. We used the default preprocessing pipeline, which includes realignment (motion estimation and correction), slice-timing correction, outlier detection, structural segmentation and normalization, de-noising with CompCor (Behzadi et al., 2007) and finally smoothing. A smoothing kernel of 6mm was applied, and denoising was done using a band-pass filter range of [0.008, 0.09] Hz.

#### 2.3.2 2.4.2 Data from Patients with Disorders of Consciousness

Data was acquired at Addenbrookes Hospital in Cambridge, UK, on a 3T Tim Trio Siemens system (Erlangen Germany). Ethical approval for testing patients was provided by the National Research Ethics Service (National Health Service, UK; LREC reference 99/391). A sample DOC patients with verifiable diagnosis were recruited from specialised long-term care centers. Consent was obtained from the patient’s legal representatives. Medication prescribed to each patient was maintained during scanning.” T1-weighted images were acquired with an MP-RAGE sequence (TR = 2300ms, TE = 2.47ms, 150 slices, 1 × 1 × 1mm^2^ resolution). Functional images, 32 slices each, were acquired using an echo planar sequence (TR = 2000 ms, TE = 30 ms, flip angle = 78 deg, 3 × 3 × 3.75mm2 resolution). Subjects were split into two groups: those who met the criteria for being in a minimally conscious state (MCS, n=10), and those who were in a vegetative state (VS, n=8).

Preprocessing was performed with SPM12 and MATLAB as described above. The first five volumes were removed to eliminate saturation effects and achieve steady state magnetization. Slice-timing and movement correction (motion estimation and correction) were performed as above, including outlier detection, structural segmentation and normalization, 6 mm FWHM Gaussian smoothing, and denoising with a band-pass filter with range [0.008, 0.09] Hz. To reduce movement-related and physiological artefacts specific to DOC patients, data underwent further de-spiking with a hyperbolic tangent squashing function. Next the CompCor technique was used to remove the first 5 principal components of the signal from the white matter and cerebrospinal fluid masks, as well as 6 motion parameters and their first order temporal derivatives and a linear de-trending term (Behzadi et al., 2007). Functional images were then bandpass filtered to remove low frequency fluctuations associated with scanner noise [0.008, 0.09] Hz.

### 2.4 Formation of Networks

After preprocessing, BOLD time-series data were extracted from each brain in CONN and the cerebral cortex was segmented into 1000 distinct ROIs, using the “Schaefer Local/Global 1000 Parcellation” (Schaefer et al., 2017)^4^. Due to the slow-convergence of Eq. 2, and the necessity of having a network with a wide enough diameter to accommodate a sufficiently wide range of box-sizes, we attempted to strike an optimal balance between network resolution and computational tractability.

For some DOC patients, there were ROI nodes which mapped to regions that had been so damaged that no detectable signal was recovered: these time-series were removed from analysis. For the MCS patients, the average number of removed nodes was 1.2 ± 1.53 nodes (≈ 0.12% of all nodes), while for the VS patients it was 5.38±7.12 nodes (≈ 0.54% of all nodes). We expect that the removal of such a comparatively small number of nodes to have a negligible effect on our overall-analysis. Every time-series *F* (*t*) was correlated against every other time-series, using the Pearson Correlation, forming a matrix *M* such that:

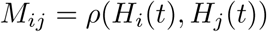

The correlation matrix has a series of ones that run down the diagonal, corresponding the correlation between each timeseries and itself which, if treated directly as a graph adjacency matrix, would produce a graph where each node had exactly one self-loop in addition to all it’s other connections. To correct for this, the matrices were filtered to remove self-loops by turning the diagonal of ones to zeros, ensuring simple graphs:

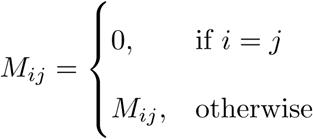

Finally, following the findings by Gallos et al., that fractal character was only present at high thresholds the matrices were binarized with a 95% threshold, such that:

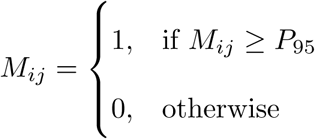

All surviving values *M*_*ij*_ < 0 ↦ 0 The results could then be treated as adjacency matrices defining functional connectivity graphs, where each row *M*_*i*_ and column *M*_*j*_ corresponds to an ROI in the initial cortical parcellation, and the connectivity between all nodes is given by Eq. 3. To see samples of the binarized adjacency matrices, see Figure 1. To see a visualization of one of the networks, see Figure 2

**Figure 1:**
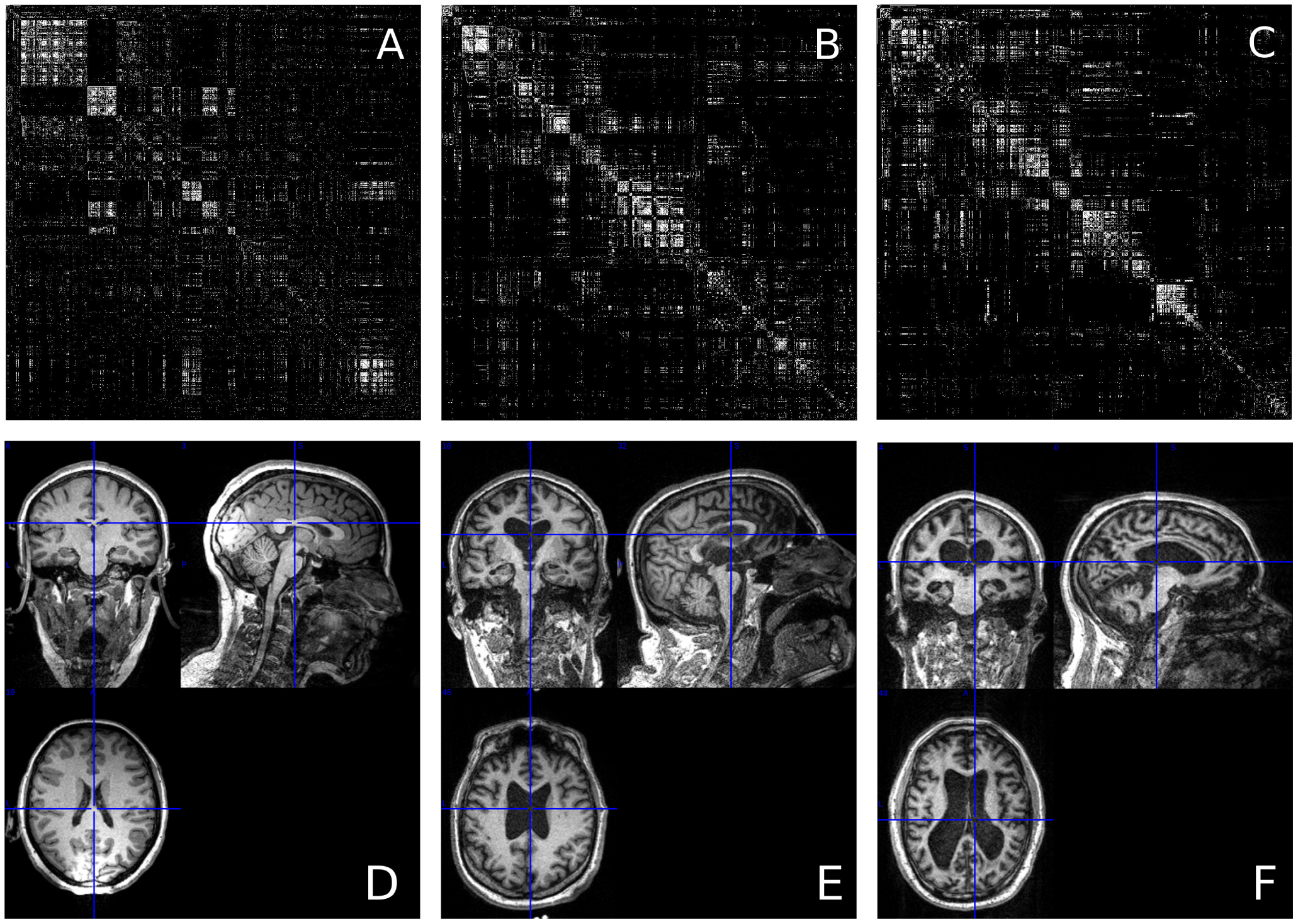
Three 1000 × 1000 adjacency matrices, representing the three conditions. *A* is a sample from the healthy control group, *B* is a sample from the MCS group, and *C* is a sample from the VS group, It is not immediately apparent that the fractal dimension of points in these groups is different. Below, see the associated structural scans (*D* is from a healthy control volunteers, *E* is from an MCS patient, and *F* from a VS patient). Note the increasingly cortical atrophy and expansion of ventricles as severity increases. Structural scans visualized in MRICron http://people.cas.sc.edu/rorden/mricron/index.html

**Figure 2:**
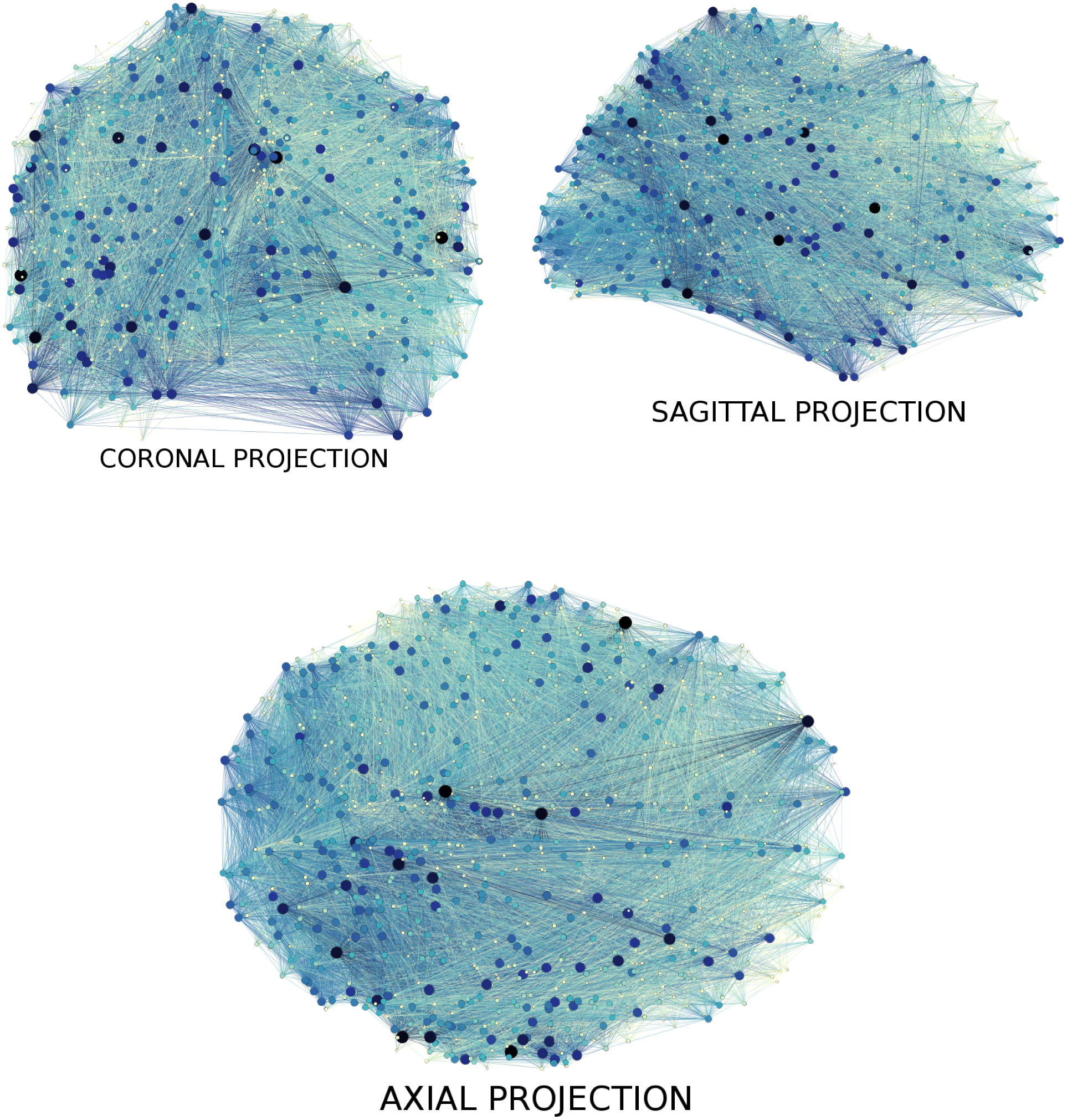
A visualization of a healthy, control functional connectivity network. Node size and darkness indicate a higher degree. Shown here are coronal, axial, and sagittal projections of the network onto a two-dimensional plane. Image made using Gephi (Bastian et al., 2009) https://gephi.org/

### 2.5 Formation of Null Graphs

To contextualize our results in the broader space of possible graphs, we generated synthetic null networks from a variety of classes to compare our three groups of functional connectivity graphs to. We generated three types of graph:

1. Lattices: a highly-ordered type of graph, where every node makes connections to it’s *k* nearest neighbours, where *k* = 2*D* and *D* is the embedding dimension of the graph. We tested two-dimensional and three dimensional lattices, each with 1000 nodes.
2. Random Graphs: A highly disordered type of graph, where every instance of the graph is selected at random from the space of all possible graphs. A population of 50 random graphs was generated and the average fractal dimension calculated. Each graph had 1000 nodes, and an identical number of edges to the natural functional connectivity networks, thresholded at 95%.

All null graphs were generated using the already-implemented graph generators in NetworkX. We hypothesized that, despite their radically different topologies, both the lattice and random graphs would have very low fractal dimensions relative to the natural functional connectivity networks when tested with the CBB algorithm.

### 2.6 Statistical Analysis

All statistical analysis was carried out using Python 3.6 using the Anaconda Python environment^5^ and Spyder IDE^6^. All packages were of the newest stable release, with the exception of the NetworkX graph analysis package (Hagberg et al., 2008): the implementation of the CBB algorithm required the use of NetworkX version 0.36. Given the heterogeneous nature, and small size, of the DOC datasets, a normal distribution was not assumed and all hypothesis tests were non-parametric. The analysis of variance was done using a one-way Kruskall-Wallis test, and then post-hoc testing was done using the Mann-Whitney U test. To control for false discoveries, p-values were tested with the Benjamini-Hochberg procedure with a false-discovery rate of 5% (Benjamini and Hochberg, 1995). All tests were from the Scipy.Stats package (Jones et al., 2001).

## 3 Results

### 3.1 Network Fractal Dimension

The Kruskal-Wallis test found significant differences between the fractal dimension of functional connectivity networks for all three conditions (H(19.91), p-value ≤ 0.0001). The median value *d*_*B*_ for the healthy control condition was 3.478 (IQR: 3.317-3.531), for MCS patients it was 3.309 (IQR: 3.21-3.438), and for VS patients it was 3.102 (IQR: 2.922-3.281). Post-hoc analysis with the MannWhitney U test found significant differences between each condition: control vs. MCS (U(13), p-value = 0.0003), control vs. VS (U(3), p-value = 0.0001), and MCS vs. VS (U(20), p-value = 0.042). For a visualization of these results see Figure 3. All p-values survived the Benjamini-Hochberg FDR correction. For a table of results see Table 1.

**Table 1:** Table of results describing the how different conditions behaved under each measures of fractal dimension. Data reported are median (IQR: 25%-75%)

| Metric | Healthy Control | MCS | VS |
| --- | --- | --- | --- |
| Network Fractal Dimension | 3.478 (IQR: 3.317-3.531) | 3.309 (IQR: 3.21-3.438) | 3.102 (IQR: 2.922-3.281) |
| Adj. Matrix Fractal Dimension | 1.731 (IQR: 1.716-1.742) | 1.706 (IQR: 1.697-1.717) | 1.693 (IQR: 1.67-1.7) |
| Temporal Fractal Dimension | N/A | 0.946 (IQR: 0.927- 0.963) | 0.912 (IQR: 0.893-0.952) |

**Figure 3:**
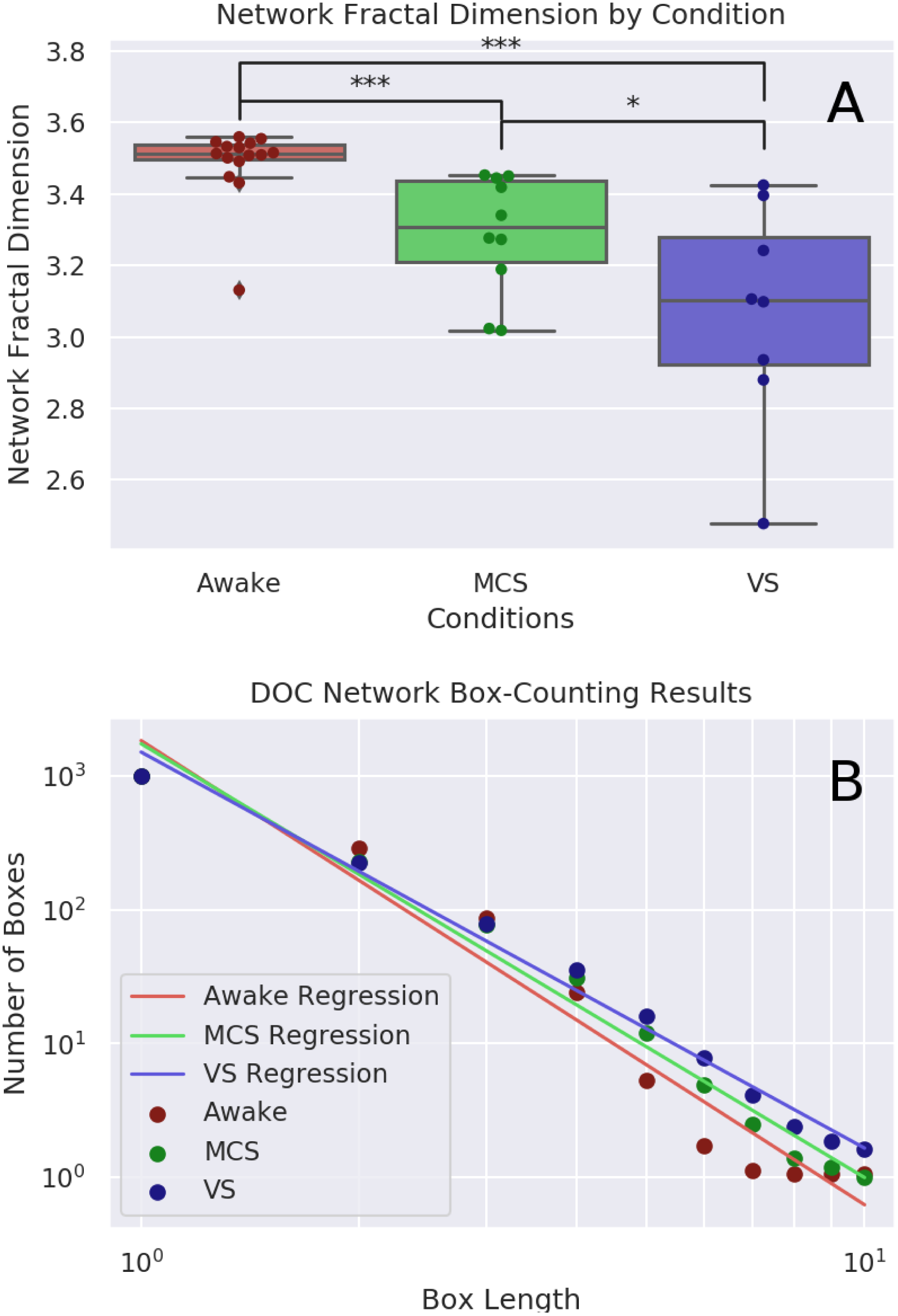
*A*: Visualization of the fractal dimension of functional connectivity networks as determined by the Compact Box Burning algorithm. Mean *d*_*B*_ for the healthy control condition: 3.37 ± 0.22 (n=15), for MCS patients: 3.29 ± 0.16 (n=10), and for VS patients: 3.07 ± 0.29 (n=8). Box length must always take integer values and does not have a regular metric unit. Post-hoc analysis with the Mann-Whitney U test found significant differences between each condition: control vs. MCS (H(41), p-value = 0.032), control vs. VS (H(23), p-value = 0.009), and MCS vs. VS (H(20), p-value = 0.042). *B* shows the relationship between *l*_*B*_ and *N* (*l*_*B*_) in all three conditions.

These results are consistent with our hypothesis that level of consciousness is positively associated with network complexity, as measured by the fractal dimension. This also shows that the direct network fractal dimension measure is sensitive enough to discriminate between different clinically useful diagnoses of grey states of consciousness, rather than simply it’s binary presence or absence.

### 3.2 Adjacency Matrix Fractal Dimension

The Kruskal-Wallis test found significant differences between the fractal dimensions of the adjacency matrices for the three conditions (H(10.24), p-value = 0.006). The median value for the healthy controls was 1.731 (IQR: 1.716-1.742), the median value for MCS patients was 1.706 (IQR: 1.697-1.717), and for VS patients it was 1.693 (IQR: 1.67-1.7). Post-hoc analysis with the Mann-Whitney U test found significant differences between the control and MCS conditions (U(33), p-value = 0.01), and the control and VS conditions (U(18), p-value = 0.0036), but not the VS and MCS conditions (U(25), p-value = 0.099). To ensure that our measures of network fractal dimension and adjacency matrix fractal dimension were associated, we correlated these values against each other and found a significant positive correlation (r=0.58, p-value = 0.0005). All the significant p-values survived Benjamini-Hochberg FDR correction. For a visualization of these results, see Figure 4.

**Figure 4:**
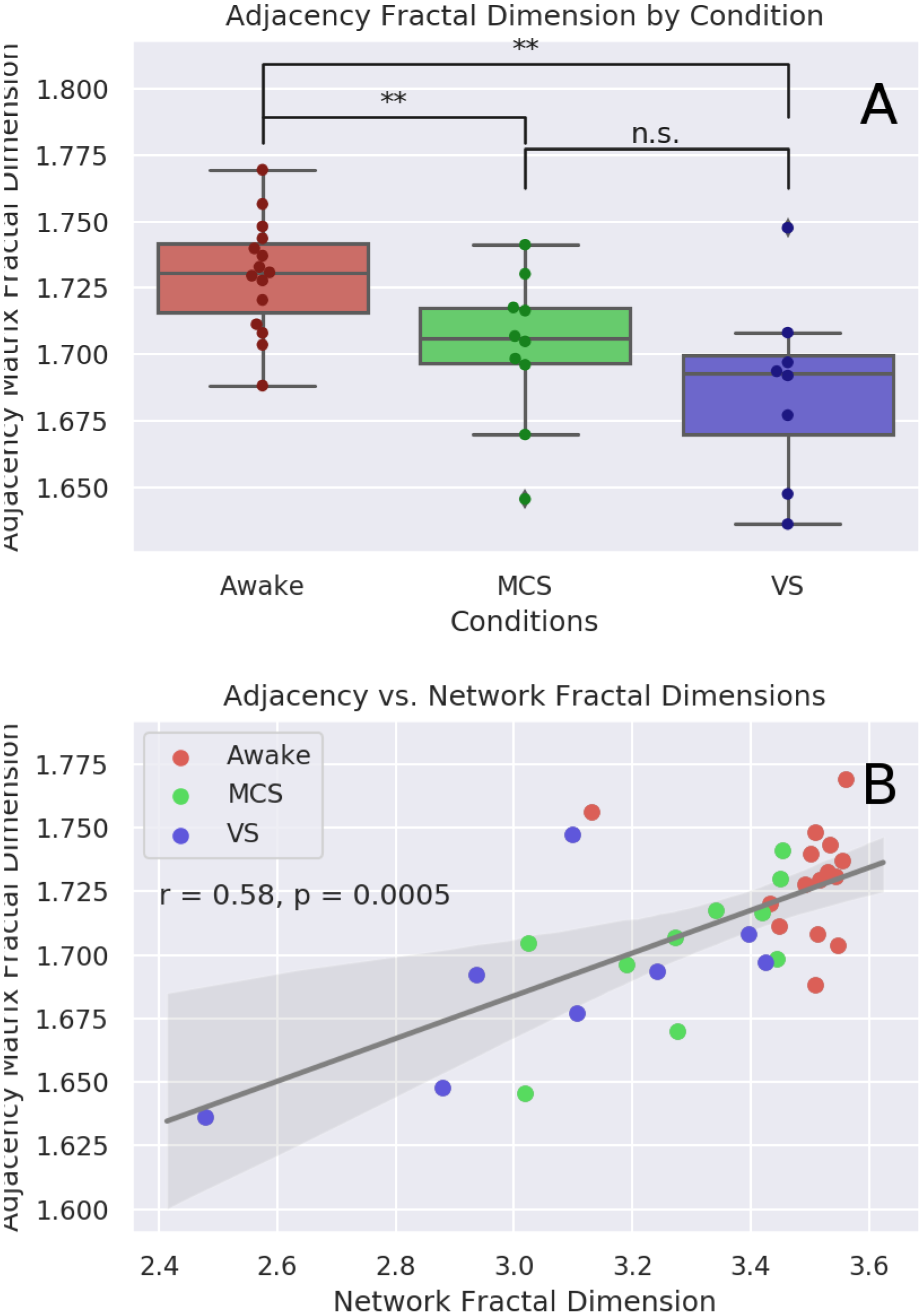
*A*: Visualization of the fractal dimension of functional connectivity networks as determined by FracLac analysis of the isomorphic two-dimensional adjacency matrix. Median value for the healthy controls: 1.731 (IQR: 1.716-1.742), for the MCS patients: 1.706 (IQR: 1.697-1.717), and for VS patients: 1.693 (IQR: 1.67-1.7). Post-hoc analysis with the Mann-Whitney U test found significant differences between the control and MCS conditions (H(33), p-value = 0.01), and the control and VS conditions (H(18), p-value = 0.0036), but not the VS and MCS conditions (H(25), p-value = 0.099). *B*: shows the correlation between the network fractal dimension (as calculated with the compact box burning algorithm), and the associated adjacency matrix fractal dimension (as calculated with FracLac).

As with the direct measure of network fractal dimension, these results show that complexity is associated with level of consciousness. While this method is sensitive enough to differentiate between healthy controls and patients with disorders of consciousness, unlike the direct measure, it was not able to discriminate between disorders of consciousness of varying severity.

### 3.3 Higuchi Temporal Fractal Dimension

Due to the large different in scan-lengths between the Awake and DOC conditions (150 samples versus 300 samples), for our analysis of temporal fractal dimension, we chose only to explore the two DOC conditions, as the Higuchi algorithm is sensitive to the length of the time-series being explored. The median value for the MCS patients was 0.946 (IQR: 0.927-0.963), and for VS patients it was 0.912 (IQR: 0.893-0.952). Testing with the Mann-Whitney U test found a significant difference between the MCS and VS conditions (U(17), p-value = 0.023). For visualization of these results, see Figure 5. We did attempt to compare all three by truncating the VS and MCS time-series to be the same length as the Awake condition, however, the resulting time-series were too short to return a meaningful answer. Surprisingly, we found no significant correlation between temporal fractal dimension and network fractal dimension or the adjacency matrix fractal dimension.

**Figure 5:**
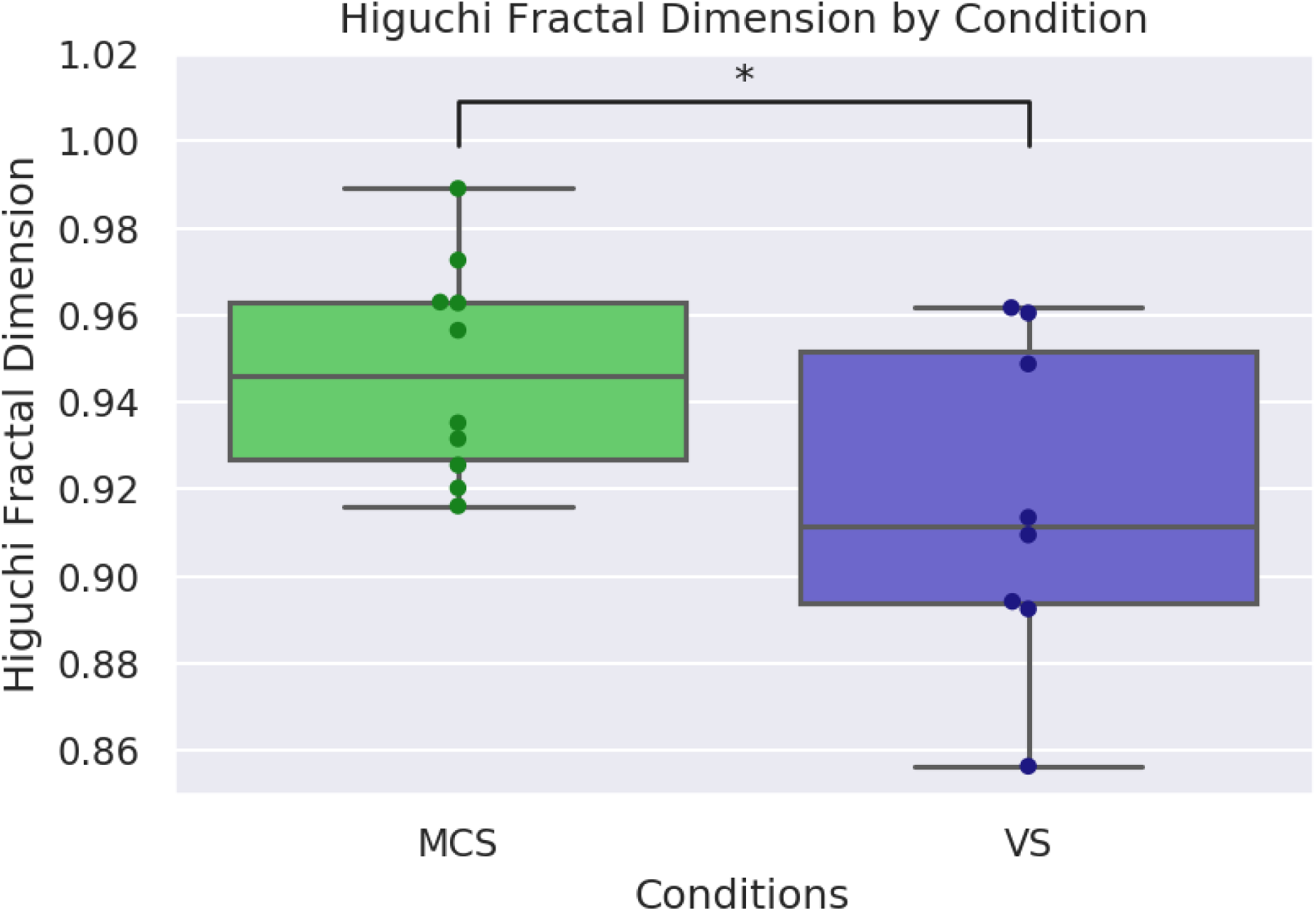
Visualization of the difference in Higuchi temporal fractal dimension between the MCS and VS conditions. As expected, the MCS condition had a higher dimension, with a median value 0.946 (IQR: 0.927-0.963) followed by the VS condition, with a median of 0.912 (IQR: 0.894-0.952). The Wilcoxon signed-rank test found a significant difference between the conditions (U(17), p-value = 0.023).

While preliminary, these results are nicely consistent with our initial hypothesis, that level of consciousness is positively associated with the fractal dimension of brain activity. Furthermore, these results complement the findings from the network fractal dimension by showing that the fractal dimension of brain activity’s relationship to consciousness is measurable in temporal, as well as spatial, dimensions.

### 3.4 Null Graph Network Fractal Dimension

As predicted, all the classes of null graphs had much lower fractal dimensions than any of our brain networks, as calculated by the CBB algorithm. The 2-dimensional lattice graph with 1000 nodes had a fractal dimension of ≈ 0.15. The 3-dimensional lattice with the same number of nodes had a fractal dimension of ≈ 0.184. The set of random networks had a higher fractal dimension, although it was still far lower than any of the real functional connectivity networks, with a median value of 0.279 (IQR: 0.279, 0.2792).

These results show that the fractal dimension measure is distinct from a measure of order/randomness, as both highly ordered networks and highly random networks return similarly low values as compared to the functional connectivity networks.

## 4 Discussion

In this study we found that the complexity of functional connectivity networks, as measured by the fractal dimension, was significantly associated with level of consciousness in healthy volunteers and patients with disorders of consciousness of varying severity. When calculated using the Compact Box-Burning (CBB) Algorithm, the fractal dimension of a these networks differentiates between healthy volunteers, patients in minimally conscious states (MCS), and patients in vegetative states (VS). A related box-counting algorithm, when applied to a two dimensional matrix isomorphic to the original graphs returned a similar result, although with less discriminative power.

The network dimension and matrix dimension results fit nicely with the previously described Entropic Brain Hypothesis (Carhart-Harris, 2018), which predicts that as the brain moves further from the zone of criticality, level of consciousness falls. If the fractal character is indicative of critical behaviour, then these results may show an association between decreased signs of criticality and disorders of consciousness. This in turn might be associated with decreased computational and information processing capabilities in the nervous system, which is turn may explain the decrease in consciousness and behavioural complexity that are the hallmarks of disorders of consciousness. While fractal dimension and entropy are distinct concepts, entropy, in computational models, positively correlates with fractal dimension (Zmeskal et al., 2013; Chen, 2016). The benefit of the fractal dimension measure, however, is that it goes beyond the order/randomness binary indexed by entropic measures (Ke, 2013).

To discuss the aetiology of the changes in network fractal dimension, we turn to previous studies, which have shown that the cerebral cortex has fractal characteristics and that changes to the fractal dimension of both the grey matter and white matter are associated with changes in cognition and the presence of clinically relevant conditions (Ha et al., 2005; Im et al., 2006; King et al., 2009; Mustafa et al., 2012). We hypothesize that the damage done to the physical cortex by brain injury translates into changes in the fractal dimension of micro-scale structural characteristics of the cortex and that this alters how individual brain regions are able to communicate. We propose that a future study that uses this same dataset to quantify changes in the fractal dimension of physical characteristics of these brains may lend evidence to this hypothesis. Specific areas of inquiry are the fractal dimension of the folds in the neocortex, which have been previously characterized as fractal, and the network of white-matter tracts revealed by DTI imaging. It would be very interesting to perform the same analysis we have reported here on a network of white-matter connections, so long as the resolution of the resulting network is high enough to support the CBB algorithm.

There are several limitations for this study that are worth considering and suggest a need for further validation. We acknowledge the comparatively small sample size, particularly in the VS condition. As previously mentioned, the requirements of fMRI image processing demand images of brains from individuals with reduced levels of consciousness, but are not so geometrically distorted as to make registration into MNI space impossible. This puts a limit on the number of brains eligible for inclusion in this kind of study. There is also the issue of parcellation resolution: we tried several different parcellations of various sizes, but only the parcellation with 1000 ROIs had a high enough resolution to return a meaningful result, and even that was still too small to permit more than 10 integer values for *l*_*B*_.

Going forward, we hope that this kind of analysis may one day be useful in a clinical context for estimating whether consciousness is present in patients who may be unable to give a voluntary behavioural affirmation of awareness. The fractal dimension measure encodes significant information about the complexity of a system into a single, easily digestible measure that seems to have predictive validity in a clinically meaningful population. As, at least in larger hospitals, MRI scans are already a routine part of clinical assessments in cases of brain damage, this measure could be incorporated into the normal course of treatment.

## 5 Conclusion

In this study, we show that high-resolution, cortical functional connectivity networks have fractal characteristics and that, in patients with disorders of consciousness induced by traumatic brain injury or anoxic brain injury, reduction in the fractal dimension is associated with more severe disorders of consciousness. This is consistent with theories that associate the content, and quality, of consciousness with the complexity of activity in the brain. Furthermore, we believe that, with refinement, this measure may inform diagnosis and stratification in a clinical setting where physicians need to make judgements about a patients consciousness in the absence of behaviourally unambiguous indicators.

## Acknowledgements

This work was supported by grants from the Wellcome Trust Clinical Research Training Fellowship to RMA (Contract grant number: 083660/Z/07/Z); the UK Medical Research Council [U.1055.01.002.00001.01 to JDP; the James S. McDonnell Foundation to JDP; the Evelyn Trust, Cambridge to JA, the National Institute for Health Research (NIHR, UK), Cambridge Biomedical Research Centre and NIHR Senior Investigator Awards to JDP and DKM; the Stephen Erskine Fellowship (Queens College, Cambridge) to EAS; the British Oxygen Professorship of the Royal College of Anaesthetists to DKM. TFV is supported by NSF-NRT grant 1735095, Interdisciplinary Training in Complex Networks and Systems. The research was also supported by the NIHR Brain Injury Healthcare Technology Co-operative based at Cambridge University Hospitals NHS Foundation Trust and University of Cambridge. We would like to thank Victoria Lupson and the staff in the Wolfson Brain Imaging Centre (WBIC) at Addenbrookes Hospital for their assistance in scanning. We would like to thank Dian Lu and Andrea Luppi for useful discussions, and all the participants for their contribution to this study.

## Footnotes

1 http://imagej.nih.gov/ij/plugins/fraclac/fraclac.html

2 SPM12; http://www.fil.ion.ucl.ac.uk/spm/

3 http://www.mathworks.co.uk/products/matlab/

4 https://github.com/ThomasYeoLab/CBIG/tree/master/stableprojects/brainparcellation/

5 https://www.anaconda.com/download

6 https://github.com/spyder-ide/spyder

